# Q1020R in the spike proteins of MERS-CoV from Arabian camels and recent human cases confers resistance against soluble human DPP4

**DOI:** 10.1101/2025.06.17.660103

**Authors:** Nianzhen Chen, Hannah Kleine-Weber, Khaled Alkharsah, Michael Winkler, Asisa Volz, Marcel A. Müller, Victor M. Corman, Christian Drosten, Markus Hoffmann, Stefan Pöhlmann

**Affiliations:** German Primate Center – Leibniz Institute for Primate Research, Göttingen, Germany; Faculty of Biology and Psychology, Georg-August University Göttingen, Göttingen, Germany; Institute of Molecular Virology, Ulm University Medical Center, 89081 Ulm, Germany; Department of Microbiology, College of Medicine, Imam Abdulrahman Bin Faisal University (IAU), Dammam, Saudi Arabia; Institute of Virology, University of Veterinary Medicine Hannover, 30559 Hannover, Germany; German Center for Infection Research, Partner Site Hannover-Braunschweig, Hannover, Germany; Institute of Virology, Charité - Universitätsmedizin Berlin, corporate member of Freie Universität Berlin, Humboldt-Universität zu Berlin, and Berlin Institute of Health, Berlin 10117, Germany; German Centre for Infection Research (Deutsches Zentrum für Infektionsforschung), partner site Charité, Berlin 10117, Germany; Labor Berlin-Charité Vivantes GmbH, Berlin 13353, Germany

**Author notes:** Contributed equally to this work Contact: Markus Hoffmann, Stefan Pöhlmann.

**Keywords:** Middle East Respiratory Syndrome, spike, DPP4, soluble DPP4

## Abstract

The Middle East Respiratory Syndrome Coronavirus (MERS-CoV) is a pre-pandemic coronavirus that is transmitted from camels, the natural reservoir, to humans and can cause severe disease. MERS cases have been documented in Arabia but not Africa, although the virus is circulating in both Arabian and African camels. Further, evidence has been provided that viruses in African camels might have a reduced capacity to cause disease. However, the underlying determinants are incompletely understood. Here, employing pseudotyped particles as model systems for MERS-CoV entry into cells, we compared cell entry of viruses from African and Arabian camels and its inhibition. We show that viruses found in Arabian camels and recent human cases are less susceptible to inhibition by human soluble DPP4 (sDPP4) than viruses from African camels, although both enter human cells efficiently and are comparably sensitive to inhibition by IFITM proteins and neutralizing antibodies. Furthermore, relative resistance to soluble DPP4 was linked to mutation Q1020R, present in the spike proteins of recent Arabian but not African viruses. These results support the concept that soluble DPP4 might constitute a natural barrier against human infection that is more efficiently overcome by viruses currently circulating in Arabian camels than those in African camels.

**IMPORTANCE:** MERS-CoV is an emerging virus that can cause severe lung disease, MERS, and is transmitted from camels to humans. Although MERS-CoV infects camels in both Africa and Arabia, MERS cases have only been documented in Arabia for reasons that remain incompletely understood. Here, we provide evidence that viruses recently circulating in Arabian camels and causing human infections are less susceptible to inhibition by human sDPP4 – a secreted version of the viral receptor that is present in various bodily fluids. Furthermore, we link sDPP4 resistance to mutation Q1020R in the spike protein of these viruses. These results suggest that viruses currently circulating in Arabian camels are better equipped to overcome a natural barrier to infection, sDPP4, than those circulating in African camels.

---

The Middle East respiratory syndrome coronavirus (MERS-CoV) is a zoonotic virus transmitted from dromedary camels to humans and causes a severe disease with a case fatality rate exceeding 30%. MERS-CoV circulates in both Arabian (clade A and B viruses) and African (clade C viruses) dromedary camel populations. However, despite serological and virological evidence of camel-to-human transmission in Africa, none of the over 2,600 confirmed human MERS cases reported to date have originated from the continent (1–3). Recent cases have been attributed primarily to clade B, lineage 5 viruses, which exhibit enhanced replicative fitness (4–6) and have predominated since 2015 (7, 8). In contrast, clade C viruses appear to be attenuated (6). Nonetheless, the molecular determinants underlying the differences in replicative capacity and virulence among MERS-CoV strains remain incompletely understood.

The MERS-CoV spike (S) protein facilitates viral entry into target cells, determines viral cell and species tropism, and is the primary target of the neutralizing antibody response. For cell entry, the receptor-binding domain (RBD) within the S1 subunit of the S protein binds to the cellular receptor, DPP4/CD26 (9–12). Subsequently, the S2 subunit facilitates fusion of the viral and cellular membranes. In order to obtain membrane fusion competence, the S protein depends on activation by host cell proteases. Upon pre-cleavage of the S protein by furin in infected cells (13–17), the serine protease TMPRSS2 cleaves and thereby activates the S protein during viral entry into cells, and its activity is essential for full viral spread and pathogenesis in animal models (18–21). Alternatively, the pH-dependent cysteine protease cathepsin L can activate the S protein in host cell endo/lysosomes (17–19), but its role in viral spread and pathogenesis is less clear.

Several factors can negatively regulate viral entry: Interferon-induced transmembrane proteins (IFITM) block cathepsin L-dependent entry of MERS-CoV (22, 23), while plasminogen activator inhibitor 1 (PAI-1) blocks TMPRSS2 activity and interferes with influenza A virus (24) and possibly also MERS-CoV proteolytic activation. Further, the cellular protein adenosine deaminase (ADA) blocks cell entry of MERS-CoV by binding to DPP4, thereby blocking S protein engagement of its receptor (25). In addition, DPP4 can be shed from the cell surface (26–28), which might require activity of metalloproteinases (29), and soluble DPP4 (sDPP4) can block MERS-CoV entry into cultured cells and viral spread in experimentally infected animals (30, 31). Finally, neutralizing antibodies from convalescent patients inhibit MERS-CoV cell entry, mainly by preventing the binding of the RBD to DPP4 (1, 32–35).

Here, we investigated whether African and Arabian MERS-CoV viruses differ in their capacity to enter cells and/or are differentially sensitive to negative regulators of the entry process. We provide evidence that viruses from Saudi Arabian camels are less sensitive to sDPP4 than their African counterparts and show that a signature mutation in the S protein of the Arabian viruses, Q1020R, contributes to this phenotype.

## RESULTS

We investigated whether the S proteins of viruses from African (n = 10) and Arabian (n = 2) camels exhibit distinct biological properties, employing a previously described cell-cell fusion assay and pseudotyped viral particles (36, 37). The S protein of MERS-CoV EMC/2012 (EMC, obtained from a human case) served as reference. The S proteins studied differed in up to 13 mutations located in several domains, including the RBD and heptad repeat 1 (HR1) (Figure 1A). All S proteins drove robust fusion of 293T effector cells with Calu-3 target cells (Figure 1B). The only exception was the S protein of Egypt-1, which was inactive due to a mutated cleavage site, as expected (38). Furthermore, all S proteins were efficiently cleaved and incorporated into particles (Figure 1C), enabling a direct comparison for host cell entry and its inhibition. Finally, all S proteins except Egypt-1 were able to drive robust entry into BHK21 cells transfected to express human and camel DPP4 (Figure 1D), consistent with expectations. In summary, all S proteins studied were efficiently expressed and were able to efficiently drive both cell-cell and virus-cell fusion. The S protein of Egypt-1 was inactive, as expected, but was included as negative control in our further analyses.

**Figure 1.**
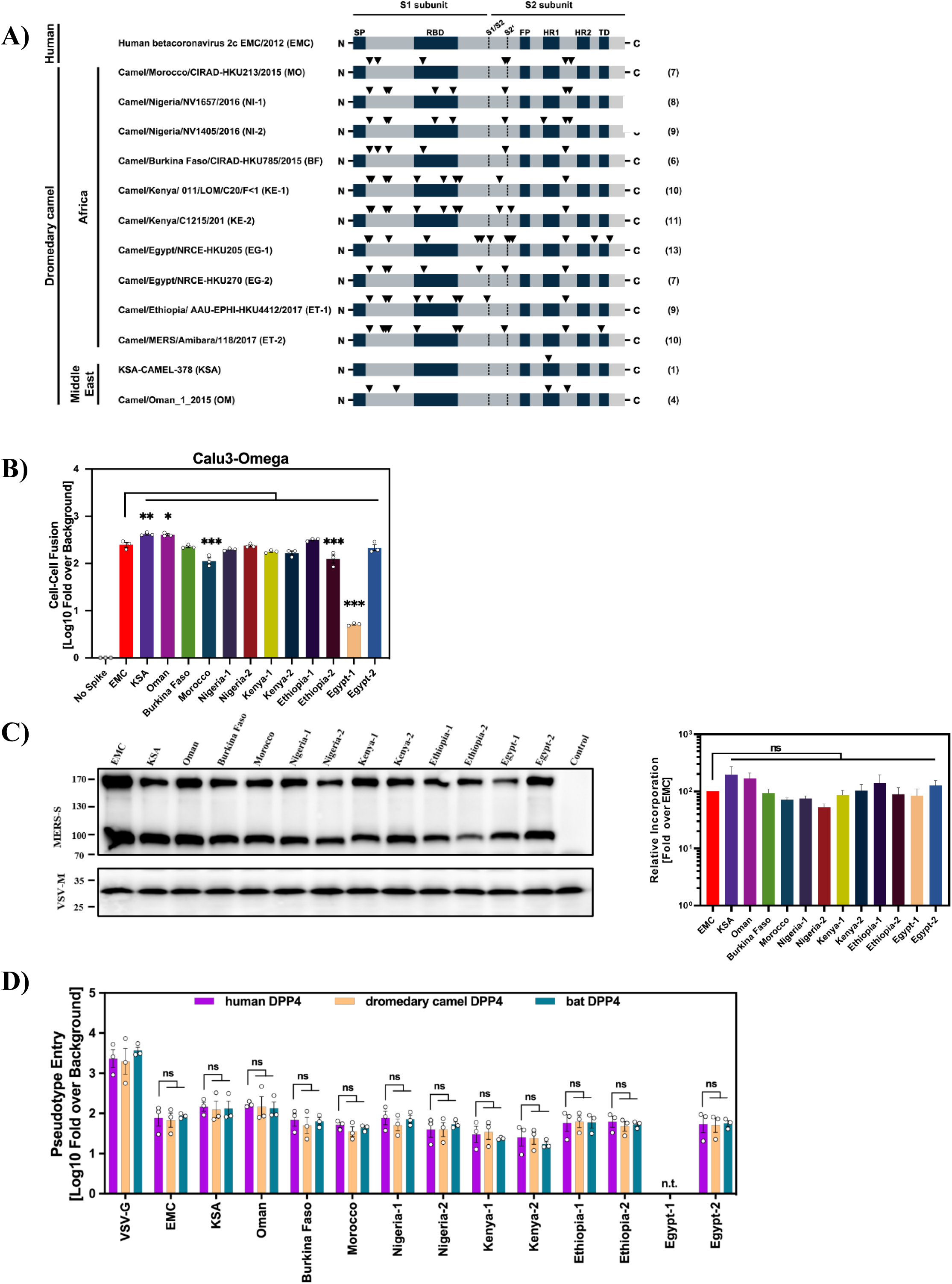
Virion incorporation, cell-cell fusion and DPP4 usage of the MERS-CoV S proteins analyzed. (A) Domain organization of the MERS-CoV S protein and mutations (black arrowheads) found in viruses from African and Arabian dromedary camels, compared to the human betacoronavirus 2c EMC/2012 (EMC) S protein. Abbreviations: SP = signal peptide, S1/S2 and S2’ = cleavage sites, RBD = receptor binding domain, FP = fusion peptide, HR1/HR2 = heptad repeat 1/2, TD = transmembrane domain. (B) For analysis of cell-cell fusion, 293T effector cells coexpressing the indicated S proteins (or no S protein) along with the beta-galactosidase alpha fragment were co-cultured with target cells (Calu-3) stably expressing the beta-galactosidase omega fragment (Calu3-Omega). At 24 h post cocultivation, beta-galactosidase activity was measured in cell lysates. The average data from three independent biological replicates are presented in panels A and B, each performed in technical quadruplicates. Data were normalized against fusion measured in the absence of S protein. Error bars indicate the SEM. Statistical significance of differences between EMC spike and the other S proteins studied was assessed by one-way analysis of variance (ANOVA) with Dunnett’s posttest (p > 0.05, not significant [ns]; p ≤ 0.05, *; p ≤ 0.01, **; p ≤ 0.001, ***). (C) For analysis of particle incorporation of the S proteins under study, vesicular stomatitis virus-based pseudotypes (VSVpp) harboring the indicated S proteins were concentrated by centrifugation and analyzed by Western Blot analysis for efficiency of S protein incorporation using an anti-V5 antibody. Detection of VSV-M served as loading control. The results of a representative blot are shown in the left panel. Right panel: Immunoblots (n = 7) were quantified using ImageJ software. Combined S protein signals (uncleaved [S0] and cleaved [S2]) were normalized against the corresponding signal of the loading control (VSV-M). Mean values are shown, error bars indicate the standard error of the mean (SEM). Statistical significance of differences in particle incorporation efficiency between human betacoronavirus 2c EMC/2012 (EMC) and the other S proteins analyzed was determined by one-way ANOVA with Dunnett’s posttest (p > 0.05, not significant [ns]; p ≤ 0.05, *; p ≤ 0.01, **; p ≤ 0.001, ***). (D) In order to analyze usage of DPP4 for cell entry, BHK-21 cells transiently expressing DPP4 of human, dromedary camel or bat origin were inoculated with pseudotypes bearing the indicated S proteins or VSV-G. At 16 h post inoculation, pseudotype entry was measured by quantifying luciferase activity in cell lysates and normalized against particles without viral envelope protein. The average of three independent experiments conducted with triplicate samples (each performed with four technical replicates) is shown. Error bars indicate the SEM. Statistical significance was tested by two-way ANOVA with Dunnett’s posttest (p > 0.05, not significant [ns]; p ≤ 0.05, *; p ≤ 0.01, **; p ≤ 0.001, ***). n.t., not tested.

### The S proteins of Arabian and African viruses mediate efficient entry into human cell lines and exhibit comparable dependence on activating proteases

We next investigated whether the robust entry into cells transfected to express DPP4 translated into efficient entry into cell lines endogenously expressing DPP4. For this, we employed 293T (kidney, human) cells stably expressing human DPP4, Huh-7 (liver, human), Vero cells (kidney, African green monkey), Caco-2 (intestine, human) and Calu-3 (lung, human) cells as targets. All S proteins except Egypt-1 mediated efficient entry into the cell lines tested (Figure 2A). Minor differences were detected but were not consistently observed among all cell lines studied. Notably, entry driven by the S proteins of the Arabian viruses KSA and Oman was generally efficient while some of the S proteins of African viruses mediated entry into Vero cells with reduced efficiency (Figure 2A). Finally, all S protein-bearing particles exhibited roughly comparable sensitivity to inhibitors of the S protein-activating host cell proteases TMPRSS2 (Camostat) and Cathepsin L (MDL 28170) (Figure 2B). The only exception were particles bearing the Nigeria-1 S protein, which showed reduced MDL 28170 sensitivity for reasons that remain unknown (Figure 2B).

**Figure 2.**
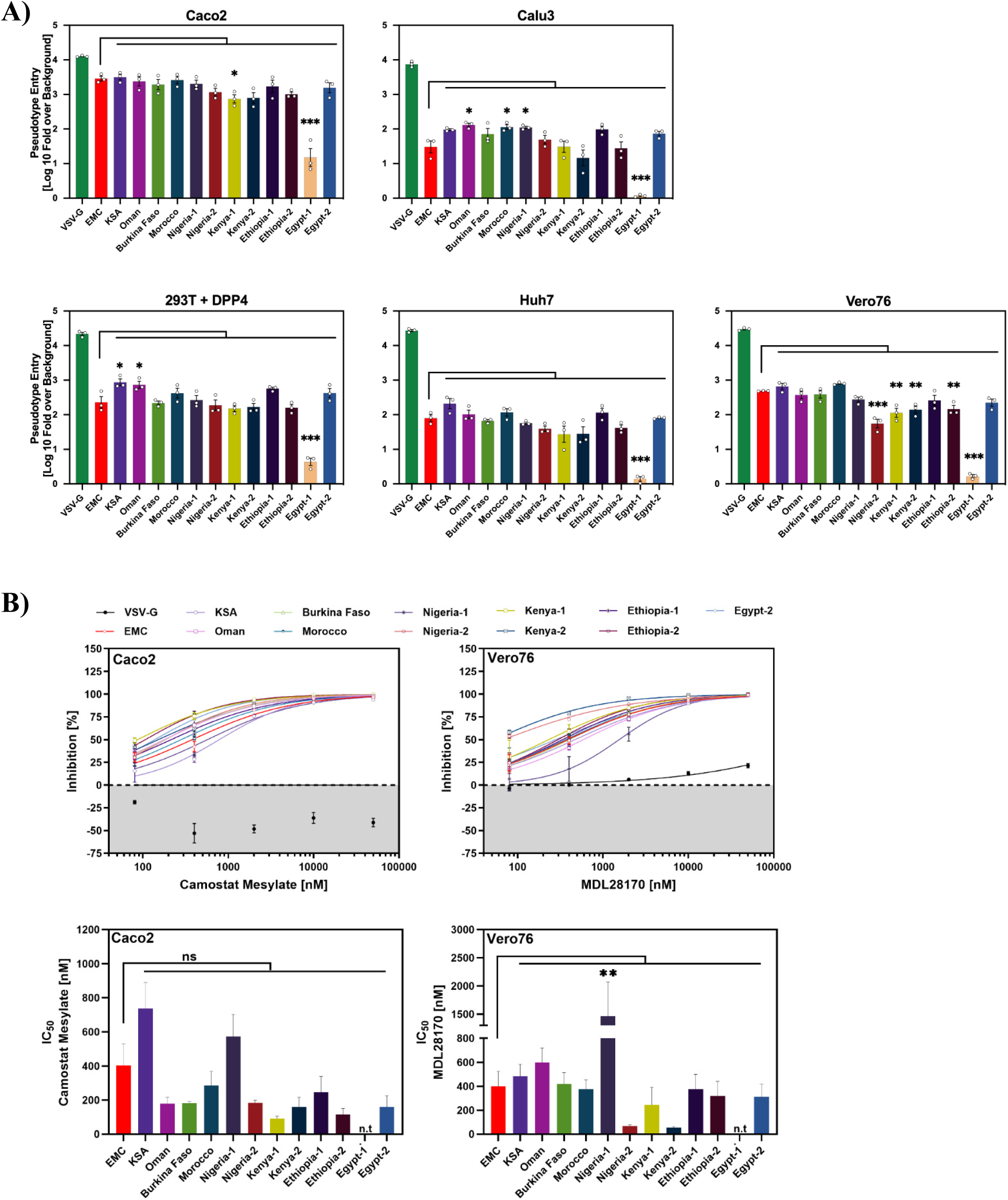
Virus-cell fusion driven by the S proteins of African and Arabian MERS-CoV is efficient and sensitive to inhibitors of cathepsin L and TMPRSS2. (A) To analyze virus-cell fusion, pseudotyped particles harboring the indicated S proteins were added onto the indicated cell lines and efficiency of cell entry was quantified by measuring the activity of pseudovirus-encoded luciferase in cell lysates at 16-20 h post transduction. The normalized average data from three independent biological replicates (each performed in technical quadruplicates) are presented, in which cell entry was normalized to particles lacking viral glycoproteins (set as 1). Error bars indicate the SEM. Statistical significance of differences between EMC spike and the other S proteins studied was assessed by one-way ANOVA with Dunnett’s posttest (p > 0.05, not significant [ns]; p ≤ 0.05, *; p ≤ 0.01, **; p ≤ 0.001, ***). (B) Protease-dependence of viral entry was analyzed by preincubating Caco-2 and Vero76 cells for 1 hour with varying concentrations (0 μM [DMSO], 0.08 μM, 0.04 μM, 2 μM, 10 μM or 50 μM) of Camostat mesylate or MDL28170, respectively, before being inoculated with pseudotyped particles harboring the indicated S proteins. At 16-18 h postinoculation, pseudovirus entry efficiency was determined by quantifying luciferase activity in cell lysates. Entry into untreated cells was set as 100% (= 0% inhibition). Data in the top panels represent the mean of three biological replicates (each performed with four technical replicates). Based on this, the inhibitor concentration leading to half-maximal inhibition of pseudovirus entry (IC_50_) was calculated and is shown in the bottom panels. Error bars represent the SEM. Statistical significance between EMC and variant was evaluated by one-way ANOVA with Dunnett’s posttest (p > 0.05, not significant [ns]; p ≤ 0.05, *; p ≤ 0.01, **; p ≤ 0.001, ***). n.t., not tested.

### Particles harboring the S proteins of African and Arabian viruses show similar temperature stability and sensitivity to IFITM proteins and neutralizing antibodies

Having established that the S proteins studied mediated robust cell entry, we next investigated the temperature stability of S protein-bearing particles and their sensitivity to IFITM proteins and neutralizing antibodies. In order to address temperature stability, the particles harboring the S proteins were incubated for different time intervals at 33, 37 and 42°C and then added to target cells, followed by quantification of entry efficiency. No apparent differences in temperature sensitivity were noted (Figure 3A). Similarly, entry of all S protein particles was comparably inhibited by stable expression of IFITM1-3 proteins in target cells, with particles harboring the hemagglutinin and neuraminidase proteins of influenza A virus A/WSN/33 (known to be IFITM-sensitive) or the glycoprotein of Machupo virus (MACV-GPC, known to be IFITM-insensitive) serving as controls (Figure 3B) (39). Finally, all S protein particles were comparably neutralized by sera from MERS-CoV infected or vaccinated and infected camels (Figure 3C and Table 1).

**Figure 3.**
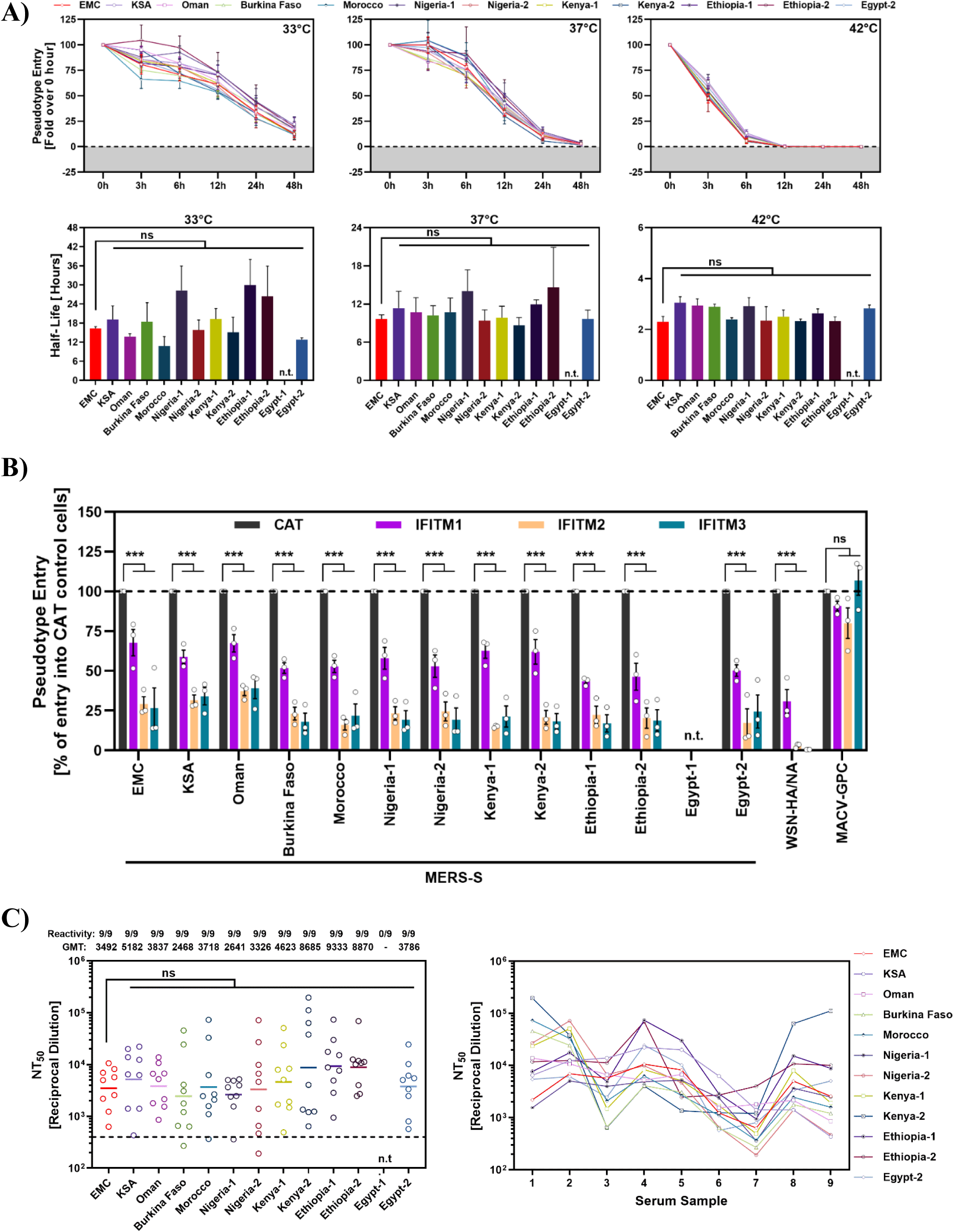
Particles pseudotyped with the S proteins of African and Arabian viruses are comparable sensitivity to temperature, IFITM proteins and neutralizing antibodies. (A) Pseudotyped particles harboring the indicated S proteins were incubated at 33 °C, 37°C, or 42 °C for the indicated durations (0, 3, 6, 12, 24, and 48 hours) before being added to Vero cells. At 16-18 h postinoculation, pseudovirus entry was measured by quantifying luciferase activity in cell lysates. Cell entry of particles pre-incubated for 0 hours at the respective temperatures was set as 100% entry. The data represent the average of three independent experiments, each conducted with triplicate samples and four technical replicates per sample. Error bars indicate the SEM. The lower panels presented the half-life derived from nonlinear curve fitting using the exponential decay model. Statistical significance was tested by one-way ANOVA with Dunnett’s posttest (p > 0.05, not significant [ns]; p ≤ 0.05, *; p ≤ 0.01, **; p ≤ 0.001, ***). n.t., not tested. (B) 293T cells stably expressing interferon-induced transmembrane proteins (IFITM1 to IFITM3) or chloramphenicol acetyltransferase (CAT; control) were transduced with pseudoviruses harboring the indicated S proteins, influenza A virus (WSN, subtype H1N1) hemagglutinin and neuraminidase (WSN-HA/NA), Machupo virus glycoprotein (MACV-GPC), or no viral glycoprotein (negative control; data not shown). At 18 h post inoculation, efficiency of pseudovirus entry was analyzed by measuring luciferase activity in cell lysates. Entry into control cells, 293T-CAT, was set as 100%. Presented are the averages from three individual experiments performed with four technical replicates. Error bars indicate the SEM. Statistical significance was tested by two-way ANOVA with Dunnett’s posttest (p > 0.05, not significant [ns]; p ≤ 0.05, *; p ≤ 0.01, **; p ≤ 0.001, ***). n.t., not tested. (C) Pseudotyped particles bearing the indicated S proteins were preincubated (30 min, 37°C) with different dilutions of plasma derived from MVA-MERS-S vaccinated (samples 1-3) or MERS-CoV seropositive (samples 4-9) camels before being inoculated onto Vero cells. At 16-18 h postinoculation, pseudovirus entry was measured by quantifying luciferase activities in cell lysates. Cell entry in the absence of plasma was set as 0% inhibition. The serum dilution leading to a 50% reduction in S protein-driven cell entry (neutralizing titer 50, NT_50_) was calculated using a non-linear regression model. Samples that yielded NT_50_ values below 1 were considered negative and manually assigned an NT_50_ value of 1. Left panel: the combined geometric mean NT_50_ values (geometric mean titers, GMT) from a single biological replicate (each conducted with four technical replicates). The proportion of samples with detectable neutralizing activity are indicated. The dashed line indicates that lowest reciprocal dilution tested. Right panel: individual NT_50_ values per serum. Statistical significance was assessed by Kruskal–Wallis analysis with Dunn’s multiple comparison test (p > 0.05, not significant [ns]; p ≤ 0.05, *; p ≤ 0.01, **; p ≤ 0.001, ***). N.t, not tested.

**Table 1.**
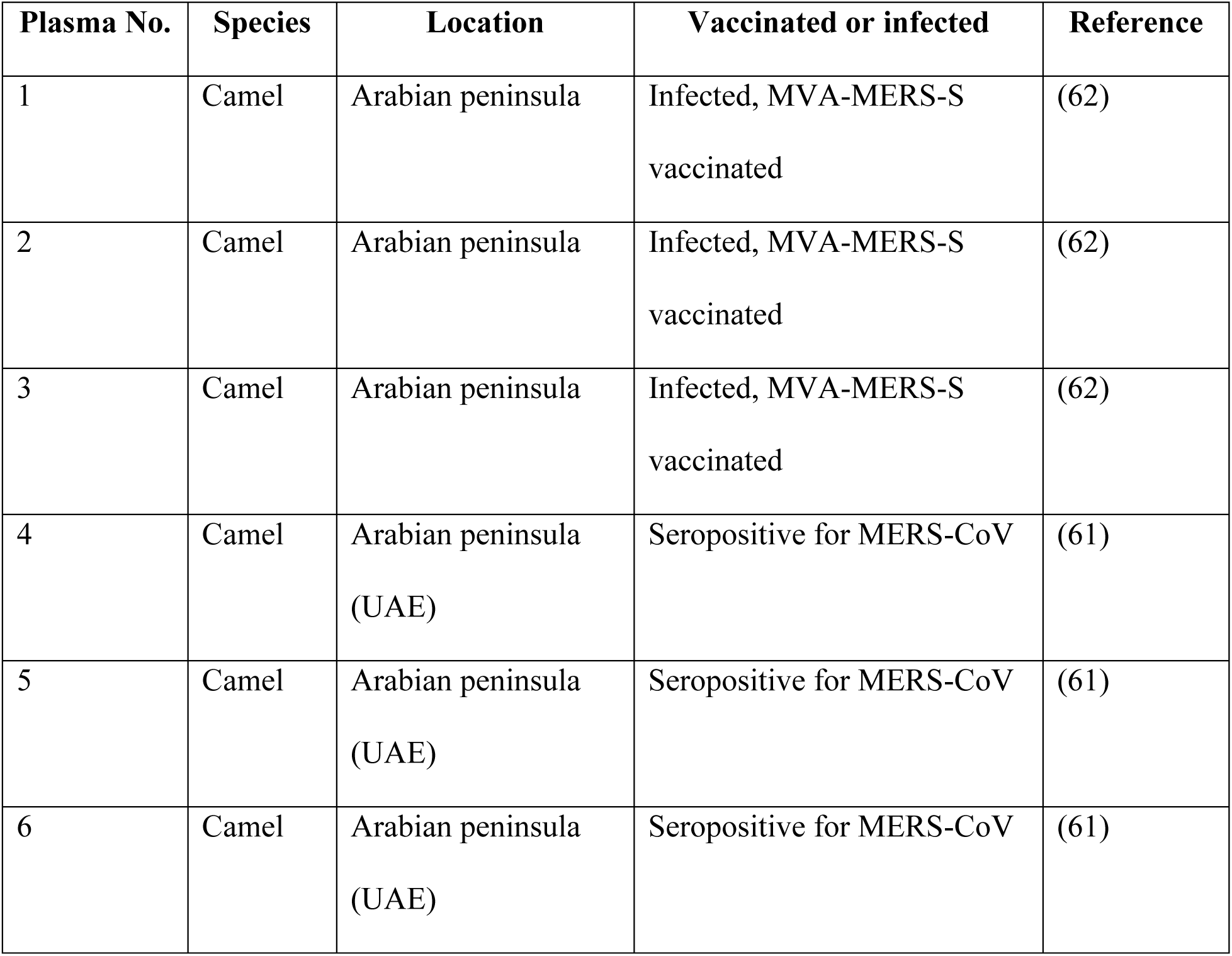
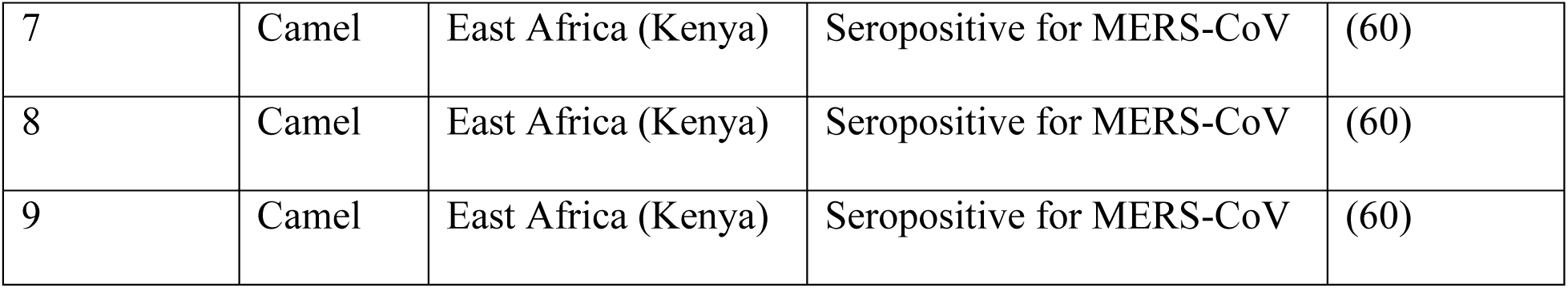

### Evidence that recent Arabian viruses are less sensitive to inhibition by soluble DPP4 due to mutation Q1020R

In humans, a soluble form of DPP4 (sDPP4) is naturally expressed in most bodily fluids (26–28), including lung fluid (40), and recombinant sDPP4 can block MERS-CoV cell entry (30, 31). Therefore, we also examined whether the S proteins studied exhibited different sensitivity to inhibition by human sDPP4. For this, we fused DPP4 to the Fc portion of immunoglobulin (sDPP4-Fc) and purified the fusion protein from 293T cells. Particles bearing the S proteins of viruses from Arabian camels were significantly less susceptible to inhibition by sDPP4-Fc as compared to particles bearing the S proteins from viruses found in African camels or EMC (Figure 4A).

**Figure 4.**
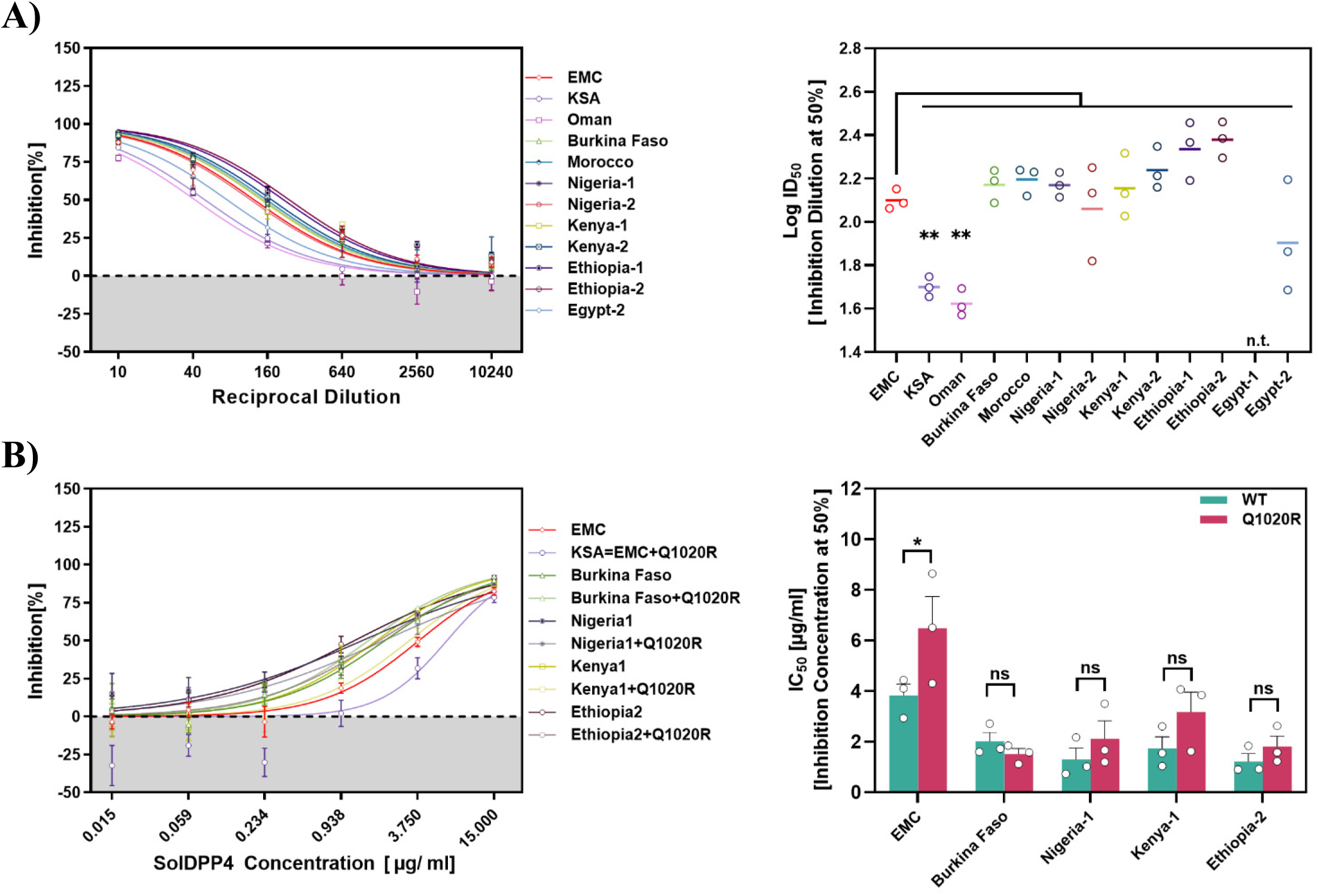
Particles pseudotyped with the S proteins Arabian viruses are less sensitive to inhibition by sDPP4-Fc and mutation Q1020R contributes to resistance. (A) Inhibition of entry by sDPP4-Fc. Particles harboring the indicated S proteins were preincubated at 37°C for 30 minutes with different dilutions of soluble human DPP4 harboring a C-terminal Fc-tag (sDPP4-Fc) before being inoculated onto Vero cells. After 16-18h inoculation, S-protein-driven cell entry was measured and normalized against particles incubated without sDPP4-Fc (= 0% inhibition). Presented are the average (mean) data ± SEM from three biological replicates (each performed with four technical replicates). The left graph shows dose-dependent inhibition of pseudovirus entry. The 50% inhibitory dilution (ID_50_) was calculated using a non-linear regression model and is presented on the right. Statistical significance of differences between particles bearing EMC S protein and the other S proteins tested was evaluated by one-way ANOVA with Dunnett’s posttest (p > 0.05, not significant [ns]; p ≤ 0.05, *; p ≤ 0.01, **; p ≤ 0.001, ***). n.t., not tested. (B) Impact of Q1020R on sDPP4-Fc sensitivity. The experiment was conducted as in panel A, but commercial sDPP4-Fc was used, S proteins with and without mutation Q1020R were analyzed and the 50% inhibitory concentration (IC_50_) and is shown in the right panel. Statistical significance between wild type and Q1020R mutation was evaluated by two-way ANOVA (p > 0.05, not significant [ns]; p ≤ 0.05, *; p ≤ 0.01, **; p ≤ 0.001, ***). n.t., not tested.

The S protein of the MERS-CoV isolate KSA-CAMEL-378, obtained from a camel from the Kingdom of Saudi Arabia (KSA), is identical at the amino acid level to S proteins of lineage 5 viruses responsible for recent human infections (8) but differs from EMC S protein only by mutation Q1020R (Figure 1A). Furthermore, this mutation was present in the S proteins of the two Arabian but not the 10 African camel viruses studied here. Therefore, we determined whether introducing Q1020R into the S proteins of EMC and selected viruses found in African camels reduced sensitivity to sDPP4-Fc. Introducing Q1020R into EMC S protein diminished inhibition by recombinant, commercial sDPP4-Fc (Figure 4B), in agreement with the data obtained with self-made sDPP4-Fc (Figure 4A). Furthermore, Q1020R reduced sDPP4-Fc sensitivity of particles bearing 3 out of the 4 S proteins from African camel viruses tested, although these effects were not statistically significant (Figure 4B).

## DISCUSSION

Our findings support the idea that sDPP4 serves as a barrier to MERS-CoV infection of humans and that the Q1020R mutation helps viruses overcome this barrier. Soluble DPP4 levels in the Saudi Arabian population are lower compared to those in other ethnic groups, and it has been suggested that this may at least in part account for the increased susceptibility to infection (41). Therefore, higher sDPP4 levels in individuals of African descent, as compared to those of Arabian descent – together with the presence of the Q1020R mutation in several Arabian but not African viruses – might contribute to the absence of MERS cases in Africa.

The IC_50_ values measured here for sDPP4-Fc-mediated inhibition of viral entry (roughly 2–6 µg/ml) are higher than the sDPP4 concentrations found in the plasma of healthy donors (roughly 0.5–0.7 µg/ml) and MERS patients (27, 31), who show reduced sDPP4 levels as compared to healthy controls (41). However, sDPP4-Fc might exert less antiviral activity than endogenous sDPP4, since dimer formation of endogenous sDPP4 is driven by non-covalent interactions between DPP4 ectodomains (42), while dimerization of sDPP4-Fc is driven by the Fc portion - likely resulting in differences in the spatial orientation of the DPP4 ectodomain. Finally, sDPP4 levels in sputum of MERS patients are reduced as compared to those in plasma (41) and levels of sDPP4 enzymatic activity in bronchoalveolar lavages (BAL) of healthy individuals are lower than those in plasma (40). However, a systematic analysis of sDPP4 levels at MERS-CoV target sites is currently missing.

Mutation Q1020R in the S protein of MERS-CoV has been detected in viruses from Arabian camels and MERS-patients and codon 1020 is under positive selection (43–46). The Q1020R mutation is located in the heptad repeat 1 (HR1) within the S2 subunit, a functional element required for S protein-driven membrane fusion. The mutation decreases stability of the 6-helix-bundle (43), the S2 conformation associated with membrane fusion. Although counterintuitive, HR destabilizing changes like Q1020R can increase infectivity in the context of other viruses (43, 47). In fact, in the present study, entry mediated by the KSA S protein was generally more efficient than that facilitated by the EMC S protein, and the KSA S protein differs from the EMC S protein only by the mutation Q1020R. Similarly, in a previous study, we found that Q1020R partially restored the reduced Calu-3 lung cell entry associated with the mutation T746K, although an impact of a second mutation, E32A, cannot be ruled out (48). Furthermore, previous studies with murine coronaviruses have revealed that residues in HR1 or other functional elements in S2 can impact receptor engagement and host range (49, 50). However, it remains to be determined how Q1020R reduces MERS-CoV susceptibility to inhibition by sDPP4.

In sum, our findings suggest that Arabian viruses harboring the Q1020R mutation are less susceptible to inhibition by human sDPP4 than viruses circulating in African camels. It will be interesting to determine in future studies whether sDPP4 is also generated in camels, exerts antiviral activity, and is differentially active against viruses bearing either Q or R at position 1020.

## MATERIAL AND METHODS

### Expression plasmids

Expression plasmids for DsRed (51), VSV-G (vesicular stomatitis virus glycoprotein) (52), MERS-S EMC (based on human betacoronavirus 2c EMC/2012 isolate) (36), Machupo virus glycoprotein (MACV-GPC, kindly provided by Michael Farzan) (53), influenza A virus strain A/WSN/33 (H1N1) hemagglutinin and neuraminidase (WSN-HA/NA) (54), human DPP4 (11), bat DPP4 (11), dromedary camel DPP4 (11) (kindly provided by Vincent Munster), and beta-galactosidase alpha and omega fragment (55) have been previously described. In addition, overlap extension PCR was utilized to generate expression vectors encoding S proteins from various regions: KSA (Kingdom of Saudi Arabia, KSA/Camel/378, GenBank: AHY22535.1), Oman (Oman, Camel/Oman_1_2015, GenBank: AQZ41296.1), Burkina Faso (Burkina Faso, camel/Burkina Faso/CIRAD-HKU785/2015, GenBank: MG923471.1), Morocco (Morocco, camel/Morocco/CIRAD-HKU213/2015, GenBank: MG923469.1), Nigeria-1 (Nigeria, camel/Nigeria/NV1657/2016, GenBank: MG923475.1), Nigeria-2 (Nigeria, camel/Nigeria/NV1405/2016, GenBank: AVN89376.1), Kenya-1 (Kenya, camel/Kenya/ 011/LOM/C20/F<1, GenBank: MK357909.1), Kenya-2 (Kenya, camel/Kenya/C1215/201, GenBank: AXP07345.1), Ethiopia-1 (Ethiopia, camel/Ethiopia/AAU-EPHI-HKU4412/2017, GenBank: MG923466.1), Ethiopia-2 (Ethiopia, camel/MERS/Amibara/118/2017, GenBank: MK564474), Egypt-1 (Egypt, camel/Egypt/NRCE-HKU205, GenBank: KJ477102.1), Egypt-2 (Egypt, camel/Egypt/NRCE-HKU270, GenBank: KJ477103.2). For all S proteins, we also constructed versions with a C-terminal V5 tag for detection by immunoblotting. The mutation Q1020R was introduced into the S protein encoding plasmids by overlap extension PCR with overlapping primers containing the mutation. The expression vector for soluble human DPP4 was constructed as follows: First, the coding sequence for the signal peptide of human CD5 (amino acid [aa] sequence: MPMGSLQPLATLYLLGMLVASCLG) was fused to the coding sequence of the Fc portion of human immunoglobulin G (IgG, aa position: 99-330, taken from the pCG1-Fc plasmid (56) that was kindly provided by Georg Herrler. Of note, the IgG sequence harbors for variations [E99D T106P, P111L, A276T] compared to GenBank: AXN93647.1) followed by the extracellular region of human DPP4 (aa position: 29-765 of NCBI Reference Sequence: NP_001366533.1) by overlap extension PCR, including a linker sequence (aa sequence CLNLQACKLDIRGR) between IgG and DPP4. Finally, the resulting sequence was inserted into the pCG1 plasmid (kindly provided by Roberto Cattaneo) making use of BamHI and XbaI restriction sites. The integrity of all PCR-amplified sequences was verified by using a commercial sequencing service (Microsynth SeqLab).

### Cell culture

The following cell lines were used in the present study: Vero cells (African green monkey kidney, female; ATCC no. CRL-1586, RRID: CVCL_0603, kindly provided by Andrea Maisner), Huh-7 cells (human liver, male; JCRB no. JCRB0403; RRID: CVCL 0336, kindly provided by Thomas Pietschmann), BHK-21 (Syrian hamster, kidney cells; ATCC no. CCL-10; RRID: CVCL_10929, kindly provided by Georg Herrler), and 293T cells (human kidney, female; DSMZ no. ACC-635; RRID: CVCL_0063) were cultured in Dulbecco’s modified Eagle medium (DMEM, PAN-Biotech) supplemented with 10% fetal bovine serum (FBS, Biochrom), 1% 100 U/ml penicillin and 0.1 mg/ml streptomycin (pen/strep) (PAN-Biotech). 293T cells stably expressing human DPP4 N-terminally linked to DsRed (293T+DPP4) (36), IFITMs (293T-IFITM1, 293T-IFITM2, 293T-IFITM3) or chloramphenicol acetyltransferase (293T-CAT) (57) were cultured under the same conditions as the parental 293T cell line, with the addition of 0.5µg/ml puromycin (Invivogen) for selection. Caco-2 cells (human colon, male; ATCC no. HTB-37; RRID: CVCL_0025) were cultured in minimum essential medium (MEM, Thermo Fisher Scientific) supplemented with 10% FBS, 1% pen/strep solution, 1% non-essential amino acid solution (PAA) and 1 mM sodium pyruvate (PAN-Biotech). Calu-3 cells (human lung, male; ATCC no. HTB-55; RRID: CVCL_0609, kindly provided by Stephan Ludwig) were cultured in DMEM/F-12 medium (Thermo Fisher Scientific) supplemented with 10% FBS, 1% pen/strep solution, 1% non-essential amino acid solution and 1 mM sodium pyruvate. Calu-3 cells stably expressing the beta-galactosidase omega fragment (Calu3-Omega) were cultured in the same medium as the parent Calu3 cells, supplemented with puromycin (0.5 µg/ml). All cell lines were maintained in a humidified incubator at 37°C with 5% CO_2_. Cell lines were validated by short-tandem-repeat analysis, amplification and sequencing of a fragment of the cytochrome c oxidase gene, microscopic examination and/or assessment of growth characteristics. In addition, all cell lines were routinely tested for mycoplasma contamination. Transfection of 293T cells was performed by calcium-phosphate precipitation.

### Production of soluble human DPP4-Fc

An expression plasmid for human soluble DPP4 (sDPP4), linked to the Fc region of human immunoglobulin G, was transfected into 293T cells after 24 h seeding. The medium was changed after 16 h and cells incubated for an additional 32 h. The supernatant was collected on the fourth day after transfection, clarified from cells and debris by centrifugation (2,000 × g, 10 min, 4 °C) and stored at 4 °C. In addition, fresh medium was added to the cells and the procedure was repeated on the fifth day after transfection and the clarified supernatants were combined. Subsequently, sDPP4 from the remaining intact cells was harvested as follows. First, a small amount of fresh culture medium was added to the cells (2 ml/T75 flask), followed by two freeze-thaw cycles (2 h 37 °C, 1 h 37 °C) to break down the cellular membranes. Next, the cellular debris was pelleted by centrifugation on and the clarified supernatant was mixed with the combined supernatants of day 4 and 5. The material was subsequently loaded onto 50 kDa MWCO (molecular weight cut-off) Vivaspin Protein Concentrator columns (Sartorius) and centrifuged (4,000 × g, 4 °C) until a 50-fold concentration factor was achieved. Next, sDPP4 was purified using NAb™ Protein A/G Spin Columns (Thermo Fisher Scientific, Cat: 89962) according to the manufacturer’s instructions. After purification, the sDPP4 was further concentrated 30-fold using Vivaspin Protein Concentrator columns (50 kDa MWCO), aliquoted, and stored at -80 °C until use.

### Production of pseudotyped particles

Vesicular stomatitis virus pseudotype particles bearing the MERS-CoV S proteins under study, influenza A virus (WSN, subtype H1N1) hemagglutinin and neuraminidase (WSN-HA/NA), Machupo virus glycoprotein (MACV-GPC), VSV-G, or no viral glycoprotein (negative control, particles were produced in cells transfected with DsRed expression plasmid) were generated using a replication-deficient VSV vector that lacks the genetic information for VSV-G and instead expresses two reporter proteins, enhanced green fluorescent protein (eGFP) and firefly luciferase (FLuc) reporter genes, VSV*ΔG-FLuc (kindly provided by Gert Zimmer) (58), using an established protocol (51). In brief, 24 h posttransfection with plasmid encoding the desired viral glycoprotein, 293T cells were infected at a multiplicity of infection (MOI) of 3 with VSV*ΔG-FLuc and incubated for 1 h at 37°C. Following this incubation, the inoculum was removed and cells were washed in phosphate-buffered saline (PBS). To neutralize any residual input virus containing VSV-G, DMEM medium supplemented with anti-VSV-G antibody (culture supernatant from I1-hybridoma cells; ATCC no. CRL-2700) was then added (of note, the antibody was not added to the medium of cells transfected with VSV-G expression plasmid). The culture supernatant was harvested and centrifuged (4,000 × g, 10 min) to remove cellular debris after 16–18 h of incubation. The clarified supernatants were subsequently aliquoted and stored at -80°C.

### Immunoblot

In order to evaluate MERS-S protein incorporation into VSV pseudotype particles, pseudoviruses expressing wild-type (EMC) or mutant MERS-S proteins with a C-terminal V5 tag were concentrated by high-speed centrifugation (13,300 rpm, 90 min, 4°C) through a sucrose cushion (20% w/v sucrose in PBS). The concentrated particles were then lysed in an equal volume of 2 x SDS-sample buffer (0.03 M Tris-HCl, 10% glycerol, 2% SDS, 5% β-mercaptoethanol, 0.2% bromophenol blue, 1 mM EDTA), heated at 96°C for 15 min, and subjected to SDS-PAGE. Proteins were transferred onto nitrocellulose membranes (Hartenstein), blocked by 5% skim milk in PBS-T (PBS containing 0.05% Tween-20) for 30 min, and incubated overnight at 4 °C with primary antibodies targeting the V5 tag (1:500, Invitrogen, Cat: R960-25), or anti-VSV-M [23H12] antibody (1:1000, Kerafast, Cat: EB0011). After incubation, the membranes were washed three times with PBS-T and subsequently incubated at room temperature for 1 hour in 5% milk solution containing horseradish peroxidase-conjugated anti-mouse secondary antibodies (1:2000, goat IgG anti-mouse IgG (H+L)-HRPO [Dianova, Cat: 115-035-045]). Following three additional washes with PBS-T, membranes were treated with a homemade chemiluminescent solution (0.1 M Tris-HCl [pH 8.6], 250 g/ml luminol, 0.1 mg/ml para-hydroxycoumaric acid, 0.3% hydrogen peroxide) and visualized using the ChemoCam imaging system with ChemoStar Professional software (Intas Science Imaging Instruments). To analyze S protein incorporation into VSV particles, protein bands were quantified using ImageJ software (version 1.53C, https://imagej.nih.gov/ij/, (59)). Total S protein signals (uncleaved, S0, and cleaved, S2) were normalized against their corresponding VSV-M signals and the resulting values were further normalized against the EMC S protein (set as 1).

### Cell-cell fusion assay

Effector 293T cells were transfected with expression plasmids for the respective S proteins (or an empty vector), along with a plasmid encoding the beta-galactosidase alpha fragment. At 24 h posttransfection, 293T cells were washed with PBS, resuspended in fresh medium and then added on top of 80% confluent Calu3-Omega cells (target cells, which stably express the beta-galactosidase omega fragment). Following a co-cultivation period of 24 h, beta-galactosidase substrate (Gal-Screen, Thermo Fisher Scientific) was added, and the cells were incubated in the dark at room temperature for 90 min. Subsequently, luminescence was measured using a Hidex Sense Plate Luminometer (Hidex).

### Analysis of spike protein-mediated cell entry

To evaluate cell tropism and host cell entry efficiency, target cells were seeded in 96-well plates and inoculated with equal volumes of pseudotype particles harboring different MERS-CoV S proteins, WSN-HA/NA, MACV-GPC, VSV-G or no viral glycoprotein (negative control). For experiments assessing the capability of S proteins to engage human, dromedary camel, or bat DPP4 orthologues as receptors, BHK-21 cells were transfected with plasmids encoding the respective DPP4 orthologues prior to infection. After 16-18 h inoculation, transduction efficiency was analyzed by measuring the activity of virus-encoded luciferase in cell lysates. The culture medium was first removed, and cells were lysed in PBS supplemented with 0.5% Triton X-100 (Carl Roth) for 30 min at room temperature. Lysates were then transferred into white 96-well plates, mixed with a commercial luciferase substrate (Beetle-Juice, PJK), and luminescence was measured using a Hidex Sense Plate Luminometer (Hidex).

To assess inhibition of viral entry by protease inhibitors, target cells (Vero and Caco2) were pre-incubated for 1 h in a medium containing different concentrations of the corresponding inhibitor, either MDL28170 (cathepsin L inhibitor, Sigma-Aldrich) or camostat mesylate (TMPRSS2 inhibitor, Sigma-Aldrich), prior to inoculation with pseudovirus particles. Medium containing the solvent (DMSO) was used as a negative control.

### Evaluation of sDPP4-mediated inhibition of S protein-driven cell entry

To analyze the inhibitory effect of sDPP4 on S protein-driven cell entry, S protein bearing particles were pre-incubated at 37°C for 30 min with different dilutions of self-made sDPP4 or different concentration of commercial sDPP4 (Biozol, Cat: BSS-BS-47084P) prior to being added to Vero cells. Particles incubated in medium without sDPP4-Fc served as control. Transduction efficiency was assessed at 16-18 h postinoculation by measuring luciferase activity in cell lysates, as described above.

### Neutralization assay

For neutralization assays, pseudovirus particles were pre-incubated (30 min, 37°C) with different serum dilutions (undiluted, 1:400, 1:1,600, 1:6,400, 1:25,600, 1:102,400) before being inoculated onto Vero cells. Of note, pseudovirus particles exposed to medium without serum served as controls. Transduction efficiency was assessed 16-18 h postinoculation by measuring luciferase activity in cell lysates, as described above.

### Temperature stability

To evaluate the thermostability of the indicated spike proteins, pseudotyped particles were pre-incubated at 33°C, 37°C, or 42°C for various durations (0, 3, 6, 12, 24, and 48 hours) before being added to Vero cells. The luciferase activity in cell lysates was measured as described above and normalized to initial entry efficiency (0 h pre-incubation).

### Plasma samples

No sampling was conducted for the present study, previously published plasma samples were analyzed (60, 61). Information on the plasma samples is provided in Table 1. All plasma samples were heat-inactivated (56 °C, 30 min) before use in neutralization experiments.

### Data analysis

Data analysis was conducted using Microsoft Excel (as part of the Microsoft Office software package, version 2016, Microsoft Corporation, Redmond, WA, USA) and GraphPad Prism version 8.3.0 (GraphPad Software, San Diego, CA, USA). Statistical analysis included one-way analysis of variance (ANOVA) with Dunnett’s posttest (incorporation efficiency, fusion assay, cell entry mediated by S protein, inhibition of entry by sDPP4), two-way ANOVA with Dunnett’s posttest (IFITM susceptibility, impact of mutation Q1020R on sDPP4 sensitivity), or Kruskal–Wallis analysis with Dunns’ multiple comparison test (neutralization assay). The serum dilutions that result in half-maximal inhibition (neutralizing titer 50, NT_50_), sDPP4 dilution factor that results in 50% inhibition (inhibitory dilution 50%, ID_50_), and the concentration of sDPP4, camostat, or MDL28170 that achieves 50% inhibition (inhibitory concentration 50%, IC_50_) were calculated using a non-linear regression model. Only p-values of 0.05 or lower were considered statistically significant (p > 0.05, not significant [ns]; p ≤ 0.05, *; p ≤ 0.01, **; p ≤ 0.001, ***).

## Acknowledgements

We thank Roberto Cattaneo, Michael Farzan, Georg Herrler, Stephan Ludwig, Andrea Maisner, Vincent Munster, Thomas Pietschmann, and Gert Zimmer for providing reagents.

